# PrioriTree: a utility for improving phylodynamic analyses in BEAST

**DOI:** 10.1101/2022.08.24.505196

**Authors:** Jiansi Gao, Michael R. May, Bruce Rannala, Brian R. Moore

## Abstract

**Summary:** Phylodynamic methods are central to studies of the geographic and demographic history of disease outbreaks. Inference under discrete-geographic phylodynamic models—which involve many parameters that must be inferred from minimal information—is inherently sensitive to our prior beliefs about the model parameters. We present an interactive utility, `PrioriTree`, to help researchers identify and accommodate prior sensitivity in discrete-geographic inferences. Specifically, `PrioriTree` provides a suite of functions to generate input files for—and summarize output from—`BEAST` analyses for performing robust Bayesian inference, data-cloning analyses, and assessing the relative and absolute fit of candidate discrete-geographic (prior) models to empirical datasets.

**Availability and Implementation:** `PrioriTree` is distributed as an R package available at https://github.com/jsigao/prioritree, with a comprehensive user manual provided at https://bookdown.org/jsigao/prioritree_manual/.

## 1 Introduction

Phylogenies are increasingly used to study the dispersal history and dynamics of pathogens. The phylodynamic methods developed by Lemey *et al*. (Lemey *et al*., 2009; Edwards *et al*., 2011) are used to infer key aspects of the geographic history of disease outbreaks, including: (1) the area in which an epidemic originated; (2) the dispersal routes by which the pathogen spread among geographic areas, and; (3) the number of dispersal events between areas.

The process of geographic dispersal among a set of discrete areas is modeled as a continuous-time Markov chain. For a geographic history with *k* areas, this stochastic process is fully specified by a *k* × *k* instantaneous-rate matrix, **Q**, where an element of the matrix, *q*_*ij*_, specifies the instantaneous rate of dispersal from area *i* to area *j*. An additional parameter, *µ*, specifies the average dispersal rate among all areas. We estimate parameters of these phylodynamic models within a Bayesian statistical framework using the Markov chain Monte Carlo (MCMC) algorithms implemented in `BEAST` (Drummond *et al*., 2012; Suchard *et al*., 2018). This approach requires that we first specify a prior probability distribution for each parameter (reflecting our beliefs about that parameter before evaluating the study data); the prior is then updated by the information in the data to return the corresponding posterior probability distribution (reflecting our updated beliefs about the parameter given our study data).

To enhance the realism of phylodynamic inferences, empirical studies frequently adopt a granular discretization of continuous geographic space into many discrete areas (Gao *et al*., 2022). However, the complexity of geographic models increases rapidly as we increase the number of areas. For example, a geographic inference problem with *k* = 5 areas has 20 parameters, with *k* = 10 areas has 90 parameters, and with *k* = 20 areas has 380 parameters. In every case, we must estimate these parameters from a dataset with minimal information; a single observation on the area where each pathogen was sampled. This inference scenario raises concerns about *prior sensitivity, i*.*e*., where posterior parameter estimates are strongly influenced by our choice of priors on the model parameters.

The inherent prior sensitivity of phylodynamic inferences is of particular concern because the priors on discrete-geographic model parameters implemented as the defaults in `BEAST`—and used in most phylodynamic studies—reflect extremely strong and biologically unrealistic assumptions about the underlying dispersal process (Gao *et al*., 2022). These considerations motivated our development of `PrioriTree`, an interactive utility for identifying and navigating prior sensitivity in discrete-geographic analyses.

## 2 Features

`PrioriTree` includes a suite of functions to generate input files for—and summarize output from—`BEAST` analyses that allow users to perform robust Bayesian inference, data-cloning analyses, and assess the relative and absolute fit of candidate discrete-geographic (prior) models to empirical datasets. We briefly outline these features below.

### 2.1 Robust Bayesian inference

`PrioriTree` allows users to assess the prior sensitivity of geographic inferences using an approach called robust Bayesian inference. This approach simply involves performing a series of MCMC analyses—of the same dataset under the same inference model—where we iteratively change one (or more) priors of our discrete-geographic model for each separate analysis. We then compare the resulting series of marginal posterior probability distributions for a given parameter to assess whether (or how much) our estimates change under different priors. While conceptually simple, the main practical challenge with robust Bayesian inference is deciding which (and how many) priors to evaluate.

`PrioriTree` allows users to explore a wide range of priors for each of the discrete-geographic model parameters. Moreover, the `PrioriTree` interface dynamically generates a graphical plot of the specified prior probability distribution to help clarify the biological implications of that prior choice. For example, an investigator may lack intuition (and/or prior knowledge) about the average rate of pathogen dispersal, *µ*; however, `PrioriTree` provides plots of the prior distribution for the number of dispersal events corresponding to a given choice of prior on *µ*. This may help guide the choice of prior; *e*.*g*., we minimally know that a pathogen that occurs in 10 areas must have experienced at least 9 dispersal events. This increased transparency may help researchers identify plausible priors to be explored via robust Bayesian analyses.

`PrioriTree` provides graphical summaries of robust Bayesian analyses, plotting distributions for a given parameter under the range of candidate priors (Fig. 1). If the inferred marginal posterior probability distributions for a given parameter are (more or less) identical under a range of corresponding priors, we can safely conclude that our estimates of this parameter are robust to the choice of prior. Conversely, if the marginal posterior probability distributions vary substantially (and resemble their corresponding marginal prior probability distributions), then we would conclude that this parameter exhibits prior sensitivity.

**Fig. 1:**
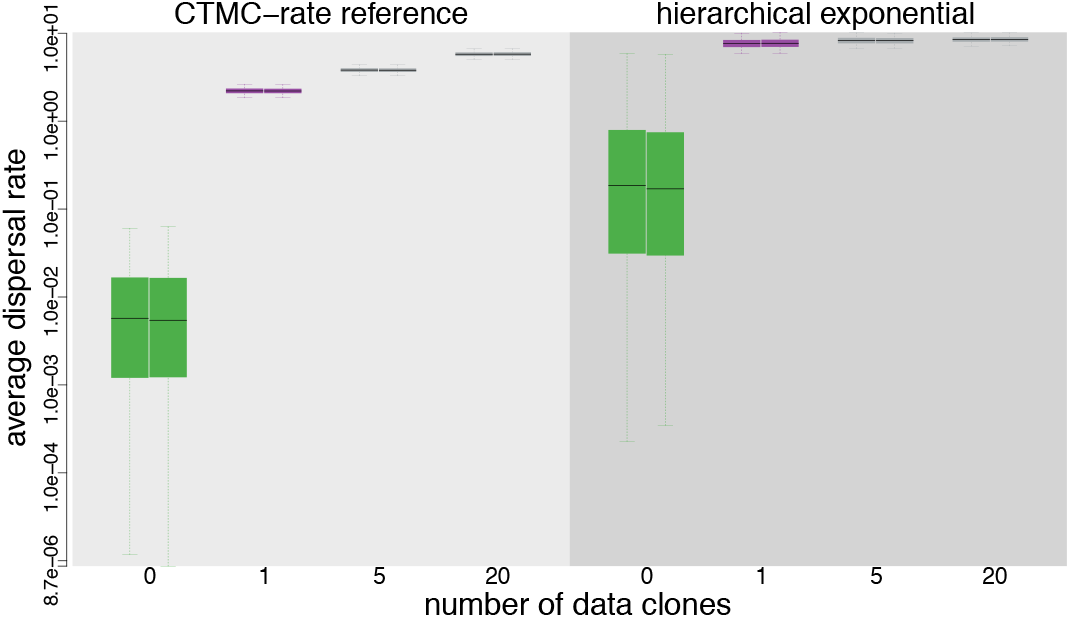
Assessing prior sensitivity in analyses of a SARS-CoV-2 dataset. PrioriTree allows users to assess the prior sensitivity of discrete-geographic analyses performed using BEAST. Here, we explore the prior sensitivity of the average dispersal rate parameter, *µ*. Panels summarize estimates of *µ* under two alternative priors; a CTMC-rate reference prior (used as the default in BEAST, left) and a hierarchical exponential prior (right). Within each panel, each boxplot summarizes the marginal distribution for *µ* inferred for different numbers of data clones (*x*-axis); the prior is inferred without data (green), the posterior is inferred from a single copy (purple), and the data-cloned posteriors are inferred from datasets with 5 or 20 copies (gray). Each pair of boxplots represents replicate analyses (to assess MCMC performance).

### 2.2 Data-cloning analyses

`PrioriTree` also allows us to assess the prior sensitivity of our discrete-geographic inferences using an approach called data cloning (Robert, 1993; Lele *et al*., 2007; Ponciano *et al*., 2009, 2012). In contrast to robust Bayesian inference, which explores prior sensitivity by assessing the impact of different prior choices, data cloning is a tool for assessing the impact of a given prior. Intuitively, data cloning measures the relative contribution of the data and the prior to the posterior distribution: an analysis is prior sensitive when the prior makes a relatively large contribution to the posterior. In practice, we perform a series of MCMC analyses—under the same inference model with identical priors—where we iteratively increment the number of copies (“clones”) of our original dataset; increasing the number of clones corresponds to increasing the relative contribution of the data to the posterior. We then explore the resulting series of posterior distributions to assess how our estimates change as the level of information in the data increases (*i*.*e*., as we increment the number of data clones).

A particular MCMC simulation in a series of data clones is defined by the number of replicate copies of our original data, *β*_*i*_. If we set *β*_*i*_ = 0, we would be targeting the joint prior probability distribution (*i*.*e*., we would be running the MCMC without data), when *β*_*i*_ = 1, we are targeting the joint posterior probability distribution (*i*.*e*., we would be running the MCMC using our original dataset). As *β*_*i*_ → *∞*, the marginal posterior distribution for the parameter under consideration will converge to a point value that is identical to the maximum-likelihood estimate (MLE) for that parameter (assuming the parameter is identifiable).

Of interest here is the relative rate at which the marginal posterior probability distribution for the parameter under scrutiny—given the prior specified for that parameter—converges to the MLE as we increase the clone number. If the prior is very informative (*i*.*e*., focused on a narrow range of parameter values) and/or the prior mean is far from the MLE value, the rate of convergence will be slow. Conversely, if a prior is more diffuse (*i*.*e*., spread over a relatively wide range of parameter values) and/or the posterior mean is relatively close to the MLE value, the rate of convergence will be relatively fast. `PrioriTree` generates summaries to assess the convergence rate by plotting marginal distributions for a given parameter under the range of *β*_*i*_ values that we explored (Fig. 1).

### 2.3 Assessing the absolute and relative fit of (prior) models

`PrioriTree` implements functions to assess the adequacy (*i*.*e*., absolute fit) of the specified (prior) model to our study data using an approach called posterior-predictive simulation (Gelman *et al*., 1996; Bollback, 2002). This Bayesian approach for assessing model adequacy is based on the following premise: if our inference model provides an adequate description of the process that gave rise to our observed data, then we should be able to use that model to simulate datasets that resemble our original data. The resemblance between the observed and simulated datasets is quantified using a summary statistic. `PrioriTree` allows users to perform posterior-predictive simulations using the output of `BEAST` discrete-geographic analyses, and then computes and plots the summary statistics to assess model adequacy (Fig. 2). `PrioriTree` also provides functions to set up power-posterior analyses for estimating the marginal likelihoods of candidate models in `BEAST` (Lartillot and Philippe, 2006; Xie *et al*., 2011; Baele *et al*., 2012) to compare the relative fit of competing (prior) models to the geographic data using Bayes factors.

**Fig. 2:**
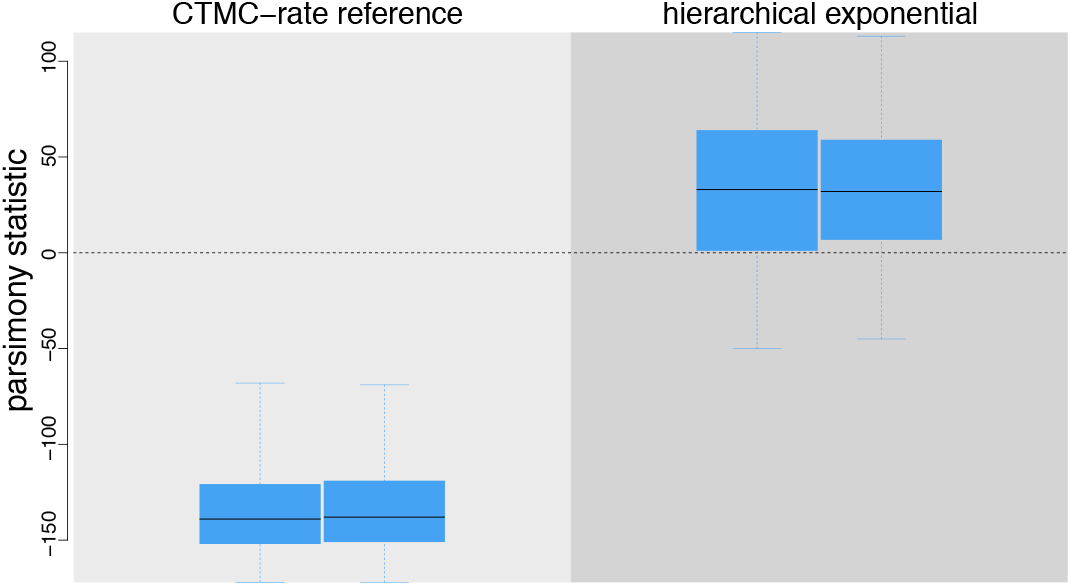
Assessing model adequacy in analyses of a SARS-CoV-2 dataset. PrioriTree allows users to assess the absolute fit of alternative prior models to the data using posterior-predictive simulation. Each boxplot depicts the posterior-predictive distribution of the summary statistic computed from datasets simulated under the a CTMC-rate reference prior on *µ* (left panel) or the alternative (hierarchical exponential) prior (right panel). Posterior-predictive distributions under the hierarchical exponential prior overlap with the observed data (dashed horizontal line), indicating that this prior model provides an adequate description of the process that gave rise to the SARS-CoV-2 dataset), whereas the CTMC-rate reference prior is inadequate. Each pair of boxplots represents posterior-predictive summaries for replicate MCMC simulations.

### 2.4 Additional Features

`PrioriTree` assumes that the phylogeny and geographic history are inferred sequentially. Under this sequential-inference approach, the phylogeny of the study group is first estimated from a sequence alignment using `BEAST`. These phylogenetic estimates are then read into `PrioriTree` as a single summary tree or as a posterior distribution of trees. If the input file contains a posterior distribution of trees, `PrioriTree` allows users to specify how to marginalize over the the distribution to accommodate phylogenetic uncertainty in the discrete-geographic inference.

Users can also set up other `BEAST` discrete-geographic inferences (*e*.*g*., inferring the number of dispersal events between each pair of geographic areas) in `PrioriTree`. In addition to generating XML scripts (as input files for `BEAST` analyses) and figures and tables (summarizing various analysis), `PrioriTree` also dynamically generates an explicit description of the methods and parameters used for each biogeographic analysis to enhance the reproducibility of phylodynamic studies.

## 3 Availability and Implementation

`PrioriTree` is developed and distributed as an R Shiny package (R Core Team, 2021; Chang *et al*., 2021)—that provides a dynamic, graphical-user interface via a local web browser—hosted on GitHub (https://github.com/jsigao/prioritree), with a comprehensive user manual available at https://bookdown.org/jsigao/prioritree_manual/.

